# Evaluating topography of mutational signatures with SigProfilerTopography

**DOI:** 10.1101/2024.01.08.574683

**Authors:** Burçak Otlu, Ludmil B. Alexandrov

## Abstract

The mutations found in a cancer genome are shaped by diverse processes, each displaying a characteristic mutational signature that may be influenced by the genome’s architecture. While prior analyses have evaluated the effect of topographical genomic features on mutational signatures, there has been no computational tool that can comprehensively examine this interplay. Here, we present SigProfilerTopography, a Python package that allows evaluating the effect of chromatin organization, histone modifications, transcription factor binding, DNA replication, and DNA transcription on the activities of different mutational processes. SigProfilerTopography elucidates the unique topographical characteristics of mutational signatures, unveiling their underlying biological and molecular mechanisms.

## BACKGROUND

Somatic mutations are found across the genomic landscapes of all cancers and of all normally functioning somatic cells [1, 2]. These mutations are carved by the activities of endogenous and exogenous mutational processes with each process exhibiting a characteristic mutational pattern, termed, *mutational signature* [3–5]. Prior studies have demonstrated that mutations are not uniformly distributed across the genome and that most mutational signatures are affected by the topographical features of the human genome [6, 7]. Specifically, mutational signatures can have distinct enrichments, depletions, or periodicities in the vicinity of early and late replicating regions [8, 9], genic and intergenic regions [10, 11], nucleosomes [12, 13], dense chromatin regions [14], histone modifications [15], and transcription factor binding sites [16, 17]. Additionally, some mutational signatures also exhibit transcription strand asymmetries, replication strand asymmetries, and/or strand-coordinated mutagenesis [18, 19].

While there is a plethora of bioinformatics tools for analysis of mutational signatures [20–32], to the best of our knowledge, only MutationalPatterns [22], TensorSignatures [31], and Mutalisk [32] consider a subset of topographical features as part of their analyses. Mutalisk performs certain topographical analysis for all somatic mutations in a sample, but it does not consider the activities of different mutational signatures which can have their own distinct topographical behaviors [32]. MutationalPatterns allows comparing the mutational patterns between different regions of the human genome and it can be used for testing enrichments or depletions using Poisson tests [22]. However, the tool does not consider the structure of the genome, the patterns of different mutational signatures, and the activities of these signatures when performing statistical comparisons. In addition, a subset of topography features has also been considered in extracting *de novo* composite mutational signatures by TensorSignatures [31], although prior benchmarking revealed sub-optimal performance when compared to traditional tools for analysis of mutational signatures [33]. In addition, the topography capabilities of all three tools are generally focused on single base substitutions and they do not support evaluating genome topography with user-provided experimental assays such as assay for transposase-accessible chromatin with sequencing (ATAC-Seq), replication sequencing (Repli-Seq), micrococcal nuclease sequencing (MNase-Seq), chromatin immunoprecipitation sequencing (ChIP-Seq), and others.

In this paper, we present SigProfilerTopography – an automated bioinformatics tool for comprehensive profiling of the topography of mutational signatures of all small mutational events, including, single base substitutions (SBSs), doublet base substitutions (DBSs), and small insertions and deletions (IDs). The tool supports examining data from a wide variety of user-provided experimental assays and can reveal dependencies between mutational signatures and chromatin accessibility, nucleosome occupancy, histone modifications, transcription factor binding sites, replication timing, transcription strand asymmetries, replication strand asymmetries, strand-coordinated mutagenesis, and other genome topography features. Moreover, SigProfilerTopography statistically compares all results with simulation data that accounts for the genome structure as well as the strengths and patterns of all operative mutational signatures within an examined sample. SigProfilerTopography is freely available for download from https://github.com/AlexandrovLab/SigProfilerTopography with an extensive documentation at https://osf.io/5unby/wiki/home/. The implementation of the tool (**Fig. 1**) and exemplars of applying SigProfilerTopography to 552 previously generated whole-genome sequenced esophageal squamous cell carcinomas (ESCCs) [34] are present in this manuscript.

**Figure 1.**
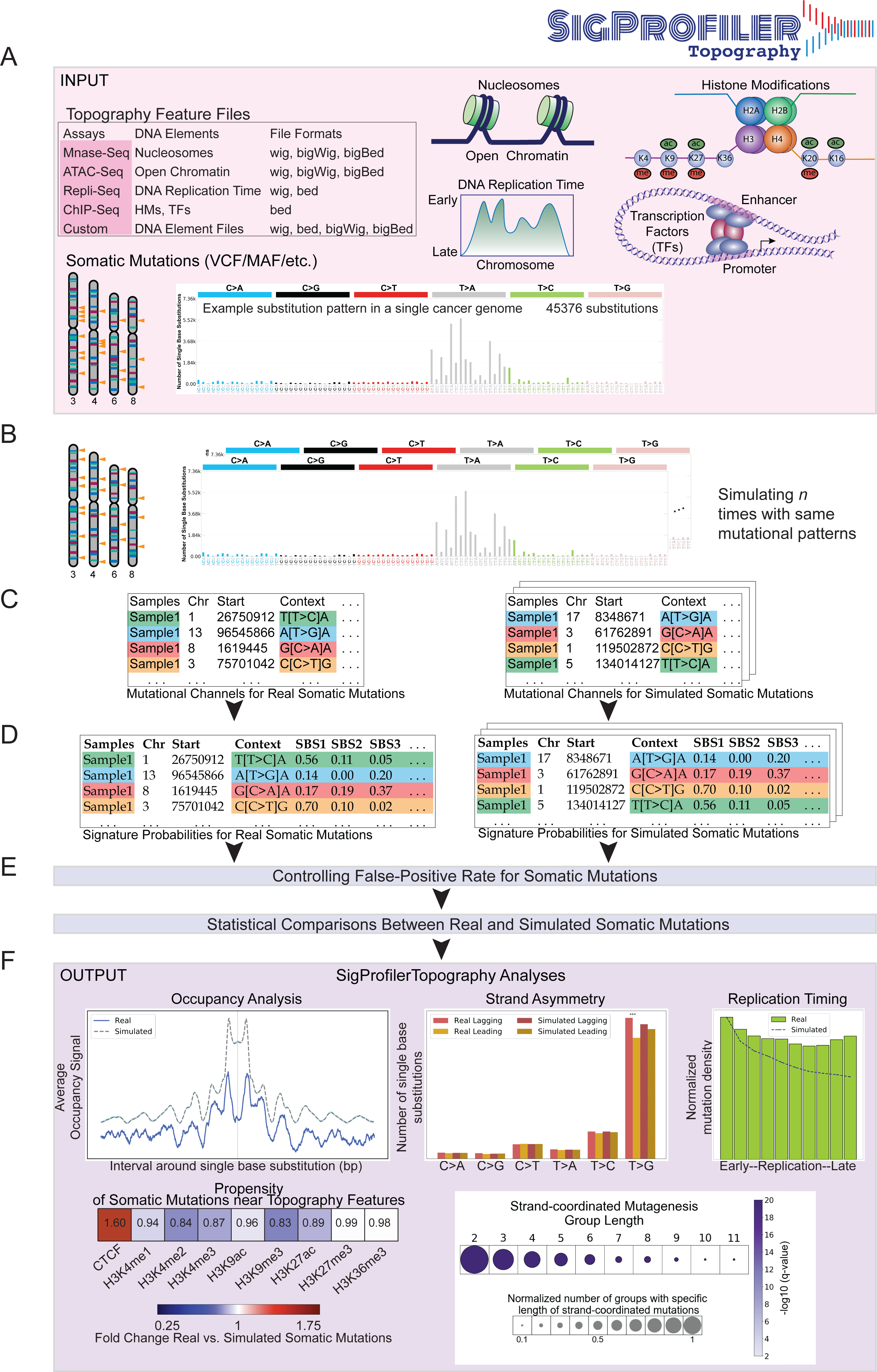
**Overview of SigProfilerTopography. *(A)*** SigProfilerTopography takes topography feature files and somatic mutations in VCF, MAF, and text formats as input. ***(B)*** SigProfilerTopography simulates real somatic mutations *n* times using SigProfilerSimulator while maintaining a preset mutational channel resolution. ***(C)*** Real and simulated mutations are annotated with mutational channel information using SigProfilerMatrixGenerator. ***(D)*** Real and simulated mutations are probabilistically attributed to different mutational signatures using SigProfilerAssignment. Alternatively, users can provide input matrices with signatures and their respective activities. ***(E)*** False-positive rates are controlled for all somatic mutations by selecting mutations highly likely to be generated by a specific mutational signature (average probability of ≥ 90% by default). For all downstream analysis, statistical comparisons are performed between real and simulated somatic mutations that are highly likely to be generated by a specific mutational signature. ***(F)*** Example outputs from occupancy, strand asymmetry, replication timing, propensity of somatic mutations near topography features, and strand-coordinated mutagenesis analyses are displayed.

## RESULTS

### Implementation and computational workflow

As input, SigProfilerTopography requires a set of topographical features of interest and a compendium of somatic mutations from a set of samples (**Fig. 1*A***). Topographical features can be derived from different genomic assays (*e.g.*, ATAC-seq, Repli-seq, MNase-seq, ChIP-seq, *etc.*) and these features can be inputted in a number of standard file formats, including: wig, bigWig, bed, or bigBed. SigProfilerTopography’s support for multiple input formats allows for topographical features to be directly downloaded from the Encyclopedia of DNA Elements (ENCODE) [35] or these features can be provided from user-generated experimental datasets. Similarly, SigProfilerTopography can examine somatic mutations using commonly supported file formats, including, Variant Call Format (VCF) and Mutation Annotation Format (MAF). By default, SigProfilerTopography utilizes SigProfilerAssignment [36] to attribute the activities of known reference mutational signatures from the Catalogue Of Somatic Mutations In Cancer (COSMIC) database [37] to each examined sample. Alternatively, if another tool for assigning mutational signatures is preferred, users can provide two additional input matrices that include the patterns and activities of all operative mutational signatures in the examined samples. In either case, SigProfilerTopography will utilize the signatures’ patterns and their activities to derive the probability for each mutational signatures to generate each type of somatic mutation [33].

After processing the input data, SigProfilerTopography simulates all somatic mutations in each sample *n* times (**Fig. 1*B***; default of *n*=100) using SigProfilerSimulator [38] while maintaining the distribution of mutations across the genome at a preset resolution (**Fig. 1*B***). By default, the preset resolution maintains the total number of mutations per chromosome and the trinucleotide pattern of each somatic mutation, which encompasses the mutated base and its immediate 5’ and 3’ base-pairs. The performed background simulations can be extensively customized depending on the appropriate scientific question [38]. Through simulating all somatic mutations, the tool generates a background model that accounts for at least a preset part of the reference genome’s structure and allows assessing any statistical differences between real and simulated somatic mutations. Both real and simulated somatic mutations are categorized in their appropriate mutation types (**Fig. 1*C***) and a mutational signature is probabilistically attributed to each somatic mutation (**Fig. 1*D***). SigProfilerTopography controls the false-discovery rate and, by default, only statistically compares mutations with an average of 90% probability of being caused by a specific mutational signature (**Fig. 1*E***). Lastly, the tool outputs a variety of results allowing to distinguish differences in the topographical distribution of real somatic mutations when compared to the distribution of simulated mutations. Example analyses include evaluations of occupancy, strand asymmetries, replication timing, enrichments/depletions, and strand-coordinated mutagenesis (**Fig. 1*F***).

### Analysis of Feature Occupancy

For a given topographical feature of interest, the tool evaluates the signal for detecting this feature in the vicinity, default of ±1 kilobase (kb) flanking regions, of each examined somatic mutation (**Fig. 2*A***). The signal is aggregated for each flaking genomic position across all somatic mutations and averaged based on all available data (**Fig. 2*A***). In the rare case of no signal being found for a specific flanking location across all mutations, the average signal is reported as zero. Occupancy analysis is jointly performed for both real and simulated somatic mutations, thus, allowing statistical comparisons of the flanking patterns and any enrichments/depletions between real and synthetic mutations. Occupancy analysis is commonly performed to evaluate the effect of nucleosome occupancy, open chromatin, transcription factor binding sites, and histone modifications on the accumulation of somatic mutations from specific mutational signatures [6, 18].

**Figure 2.**
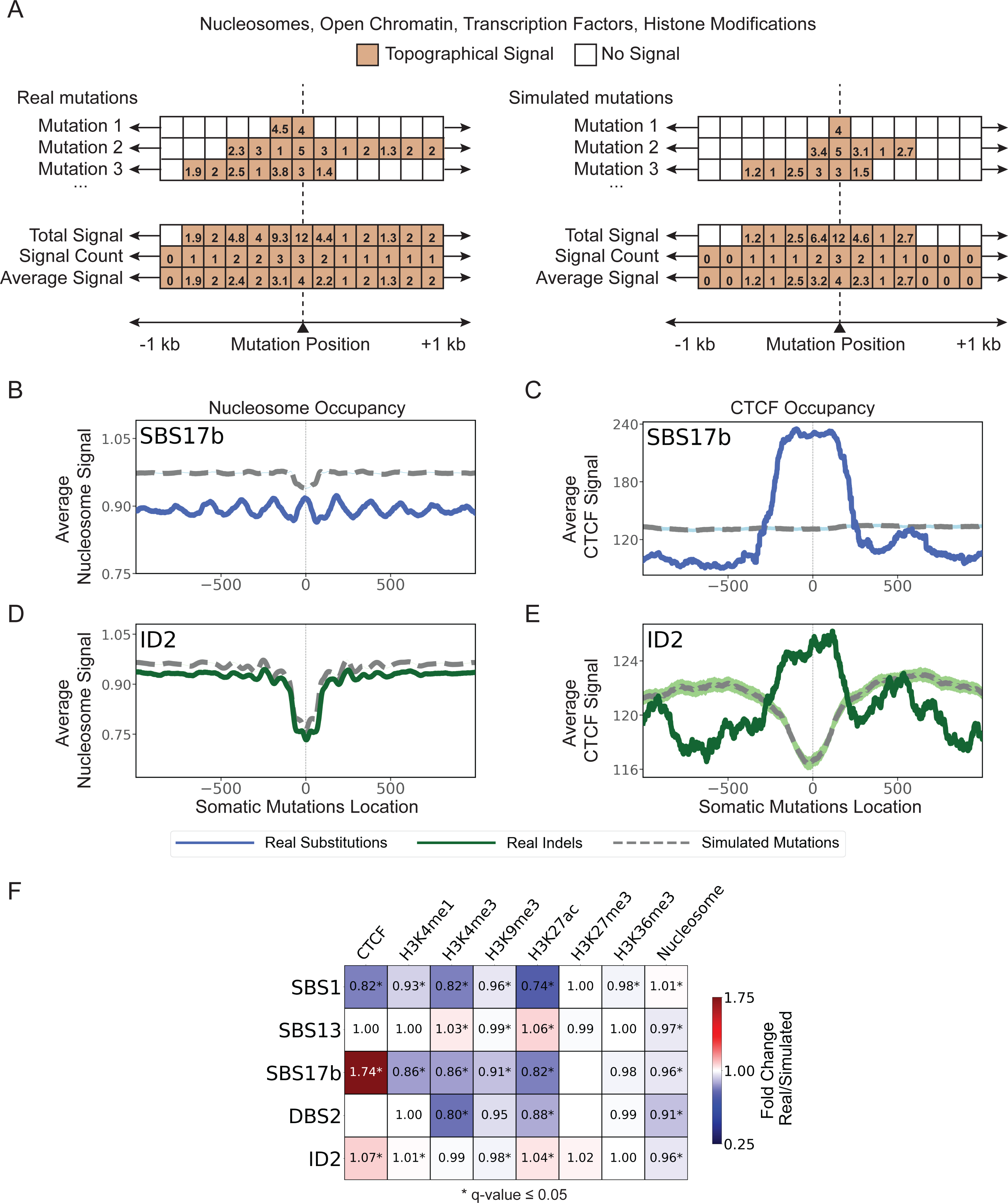
**Evaluating occupancy of topographical features. *(A)*** Conceptual and simplified depiction of SigProfilerTopography’s occupancy analysis, where x-axes correspond to ±1 kilobase (kb) from the genomic positions of real and simulated mutations. Colored boxes reflect the experimental signal detected for a specific genomic location while white boxes correspond to no experimental signal. ***(B)*** Nucleosome occupancy analysis exemplar for substitution signature SBS17b. ***(C)*** CTCF occupancy analysis exemplar for substitution signature SBS17b. ***(D)*** Nucleosome occupancy analysis exemplar for indel signature ID2. ***(E)*** CTCF occupancy analysis exemplar for indel signature ID2. In panels *(B)* through *(E)*, solid lines and dashed lines display the average topography feature’s signal (y-axes) along a 2 kilobase window (x-axes) centered at the somatic mutation locations for real and simulated mutations, respectively. The mutation location is annotated in the middle of each plot and denoted as 0. The 2 kilobase window encompasses 1,000 base-pairs 5’ adjacent to each mutation as well as 1,000 base-pairs 3’ adjacent to each mutation. ***(F)*** Heatmap displays enrichments and depletions of ESSC signatures within CTCF transcription factor binding sites, histone modifications, and nucleosomes. Red colours correspond to enrichments of real mutations and blue colours correspond to depletions of real mutations when compared to simulated data. The intensities of the red and blue colours reflect the degree of enrichments or depletions based on the average fold change. White colour boxes with no annotation correspond to insufficient data for performing statistical comparisons. Statistically significant enrichments and depletions are annotated with * (q-value ≤ 0.05).

To illustrate SigProfilerTopography’s capabilities for occupancy analysis, we examined the effect of nucleosome occupancy (measured by MNase-seq data) and binding of CTCF (based on ChIP-seq data), a key regulator of chromatin architecture, on mutational signatures SBS17b and ID2 in the ESCC cohort. Signature SBS17b has a generally unknown etiology with prior studies reporting associations with damage from reactive oxygen species [39] and possible exposure to 5- fluorouracil chemotherapy [40]. Mutations due to SBS17b exhibited periodicity with a period of approximately 190 base-pairs reflecting the nucleosome positions (**Fig. 2*B***). This periodicity has been previously attributed to high damage [41] and less repair at nucleosome positions [42]. Additionally, SBS17b substitutions were highly enriched at CTCF binding sites, which is strikingly different when compared to expected by chance from the simulated substitutions (**Fig. 2*C***). Signature ID2 has been previously attributed to slippage during DNA replication of the DNA template strand and this signature can be highly enriched in cells that are mismatch repair deficient [5]. Mutations due to ID2 were preferentially depleted at nucleosome-occupied regions (**Fig. 2*D***) while significantly enriched at CTCF binding sites (**Fig. 2*E***).

In addition to evaluating the patterns in the vicinity of a topographical feature, SigProfilerTopography allows summarizing the different enrichments and depletions of topographical features in the vicinity of somatic mutations when compared to synthetic mutations. Specifically, the tool performs a statistical test to evaluate whether the topographical signal is enriched, depleted, or as expected based on the simulated data. Applying SigProfilerTopography to 8 topographical features and 5 mutational signatures in the ESCC cohort reveals that mutational signatures can be distinctly affected by each topographical feature. For example, SBS17b is enriched in CTCF binding sites and depleted at histone marks (**Fig. 2*F***). This depletion is especially profound at H3K4me1 and H3K27ac, both of which delineate enhancer regulatory regions [43, 44].

### Evaluating Replication Timing

Cells replicate their DNA following a predefined replication timing program [45–47]. DNA replication begins simultaneously at multiple origins of replication and propagates bidirectionally on both strands. Chromosomal regions close to the origin of replication will replicate early, whereas regions that are far from the origin will replicate late. SigProfilerTopography can infer early and late replicating regions based on Repli-seq assay. Since higher signal in Repli-seq data reflects earlier replication [48, 49], the tool performs a search for local minima and maxima of the provided signal (**Fig. 3**). Specifically, weighted average data are smoothed and transformed into wavelet-smoothed signal, which results in regions with high signal values indicating domains of early replication where initiation occurs earlier in S-phase or early in a higher proportion of cells. Local maxima and local minima in the wavelet-smoothed signal data correspond to replication initiation zones (peaks) and replication termination zones (valleys), respectively (**Fig. 3*A***). SigProfilerTopography uses wavelet-smoothed signal data in replication timing analysis and, additionally, peaks and valleys data in replicational strand asymmetry analysis. After sorting the replication time signals into descending order from early to late, the tool splits the signal into deciles, with each decile containing 10% of the replication time signals. To demonstrate SigProfilerTopography’s capabilities for replication timing analysis, we evaluated the effect of replication timing in the ESCC cohort on signature SBS2 (**Fig. 3*C***), a mutational signature previously attributed to the activity of the APOBEC family of deaminases [34]. Similar to prior reports [6, 18], SBS2 exhibited an increasing normalized mutation density from early to late replicating regions (**Fig. 3*D***).

**Figure 3.**
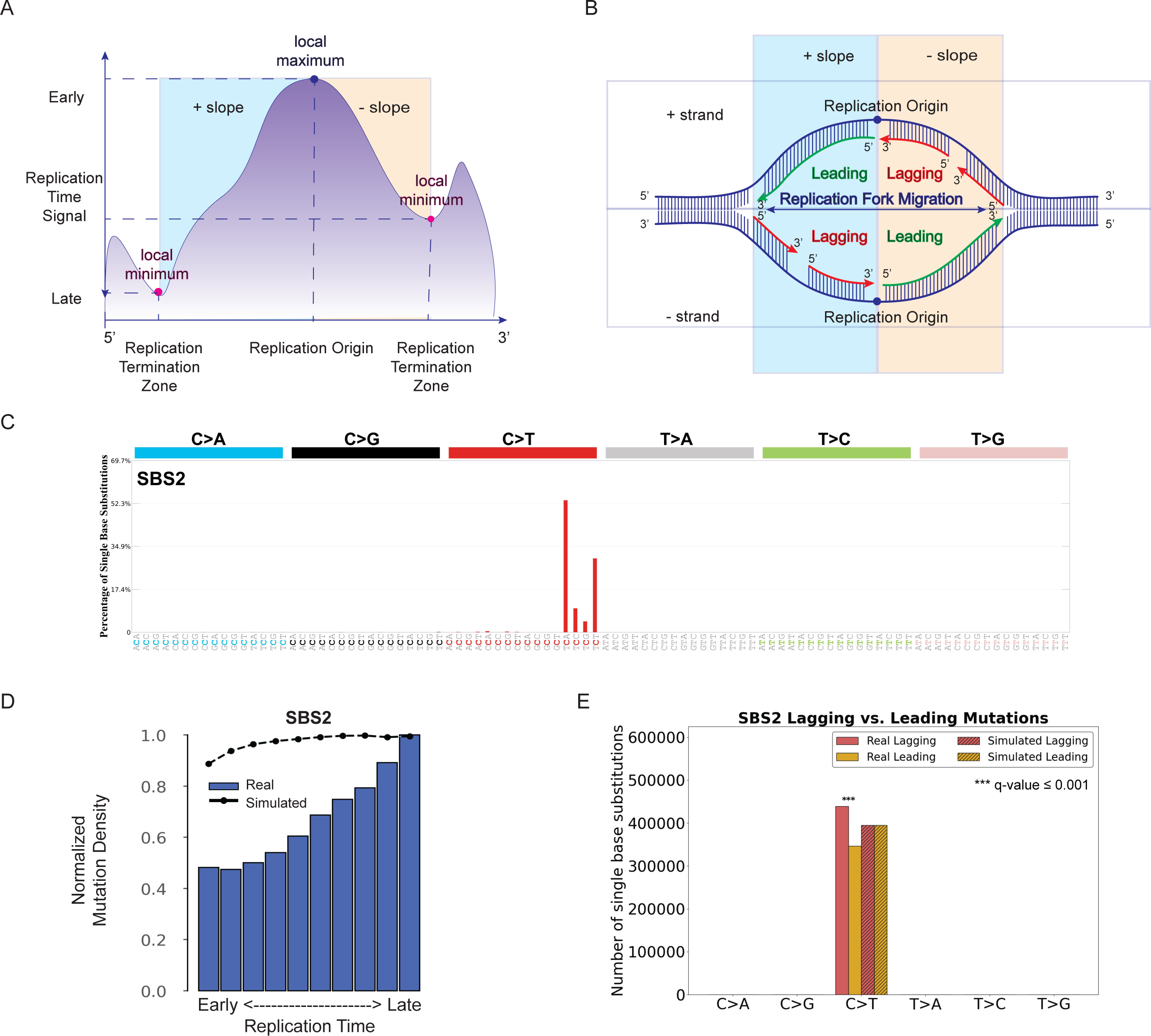
**Examining the effect of replication timing and replication strands. *(A)*** DNA replication starts at multiple origins simultaneously. Genomic regions close to replication initiation zones are replicated early, whereas genomic regions close to replication termination zones are replicated late. **(*B*)** Replicational strand classification. DNA replication starts at multiple origins of replication at the same time bidirectionally at both strands. Having the same direction for DNA synthesis and replication fork migration enables continuous DNA synthesis, which results in regions on the leading strand, whereas opposite directions of DNA synthesis and replication fork cause discontinuous DNA synthesis in small fragments, termed, Okazaki fragments, on the lagging strand. ***(C)*** Mutational profile of APOBEC-associated substitution signature SBS2 using the conventional 96 mutation type classification. ***(D)*** Replication timing analysis for substitution signature SBS2. The x-axis depicts the 10 bins from early to late replication regions, while the y- axis shows the normalized mutation density for each replication domain. The dashed line reflects the behavior of simulated mutations. **(*E*)** Replicational strand asymmetry for substitution signature SBS2. In replication strand asymmetry figure, x-axis displays six substitution subtypes based on the mutated pyrimidine base: C>A, C>G, C>T, T>A, T>C, and T>G. Mutations were oriented by the pyrimidine base of the reference Watson-Crick base-pair and classified as ones occurring on the leading or lagging strand. The y-axis represents the number of mutations on leading and lagging strands. Real and simulated mutations are shown in bar plots and shaded bar plots, respectively. Statistically significant replication strand asymmetries are depicted with * (q-value ≤ 0.05).

### Examining Replication Strand Asymmetries

In eukaryotic cells, DNA replication is initiated around multiple replication origins, from where it proceeds in both directions on both strands (**Fig. 3*B***). The strand where the direction of DNA synthesis and growing replication fork are the same is replicated continuously and it is termed leading strand. Conversely, when the direction of DNA polymerase and the growing replication fork are opposite, then that strand (termed, lagging strand) is replicated discontinuously in short Okazaki fragments [50]. Imbalance between DNA damage and DNA repair may lead to mutations from the same type to be enriched on the leading or lagging strands.

Using data from an Repli-seq assay, SigProfilerTopography can annotate mutations as ones occurring on the leading or lagging strand by orienting them by the pyrimidine base of the reference Watson-Crick base-pair. Applying SigProfilerTopography to the mutations attributed to the APOBEC-associated signature SBS2 in the ESCC cohort reveals an enrichment of mutations on the lagging strand when compared to simulated data (**Fig. 3*E***). This result is consistent with prior reports of APOBEC deaminases targeting single-stranded DNA during replication [51].

### Examining Transcription Strand Asymmetries

In addition to evaluating the effect of replication on the accumulation of mutational signatures (**Fig. 3**), SigProfilerTopography also allows examining the impact of transcription on somatic mutagenesis. Specifically, the tool annotates each mutation as either genic or intergenic, where genic mutations are within the genomic regions of well-annotated protein coding genes and intergenic mutations are outside these regions (**Fig. 4*A***). Moreover, somatic mutations within well-annotated protein coding genes are further subclassified based on the pyrimidine base of the reference Watson-Crick base-pair resulting into two additional subclasses: un-transcribed mutations and transcribed mutations (**Fig. 4*A***). This subclassification allows measuring transcription strand asymmetries due to either transcription-coupled DNA repair [52, 53] or transcription-coupled DNA damage [19]. Applying SigProfilerTopography to the somatic mutations due to SBS16 (**Fig. 4*B***), a mutational signature previously associated with alcohol consumption [54], revealed both accumulation of higher number of T>C mutations on the transcribed strand (**Fig. 4*C***) as well as an enrichment of mutations within genic regions (**Fig. 4*D***). This topographical behavior of signature SBS16 has been previously attributed to the role of transcription-coupled damage in actively transcribed genes [19, 55].

**Figure 4.**
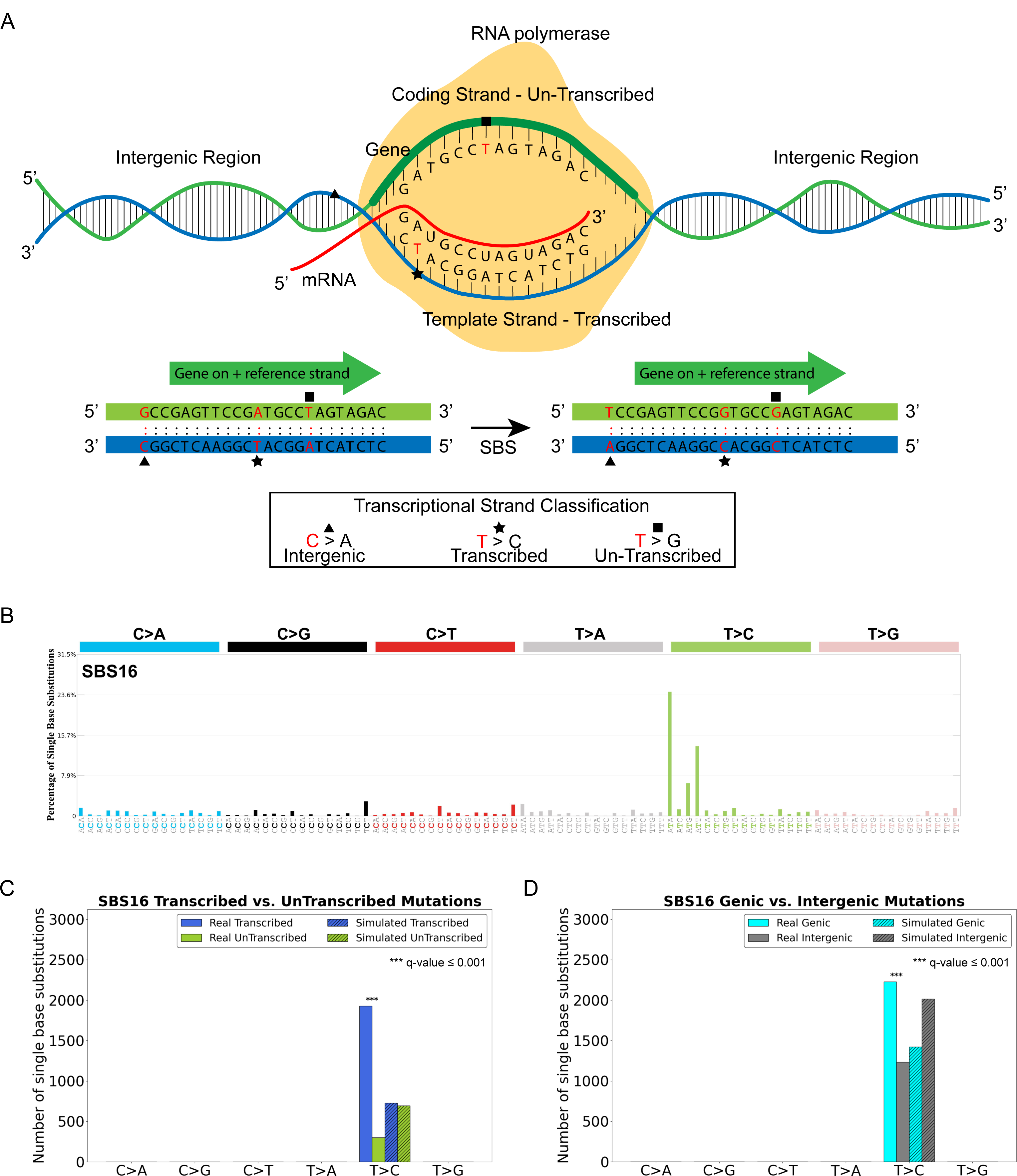
**Assessing the impact of the transcriptional machinery. *(A)*** Somatic mutations within protein coding genes are oriented by the pyrimidine base of the reference Watson-Crick base-pair and classified as ones being on the transcribed or un-transcribed strand. Somatic mutations outside protein coding genes are classified as ones in intergenic region. ***(B)*** Mutational profile of substitution signature SBS16 using the conventional 96 mutation type classification. ***(C)*** Exemplar transcriptional strand asymmetry analysis for substitution signature SBS16. X-axis displays six substitution subtypes based on the mutated pyrimidine base: C>A, C>G, C>T, T>A, T>C, and T>G, and the y-axis represents the number of mutations both for real and simulated mutations on transcribed and un-transcribed strands in bar plots. Simulated mutations are shown in shaded bar plots. ***(D)*** Exemplar genic versus intergenic regions analyses for substitution signature SBS16. X-axis is presented in a format similar to the one in *(C)*. The y-axis represents the number of mutations on genic and intergenic regions as bar plots. Simulated mutations are shown in shaded bar plots.

### Mapping Strand-coordinated Mutagenesis

Prior studies have shown that strand-coordinated mutations are commonly observed, for example, due to damage on single-stranded DNA, and can form hypermutable genomic regions [56, 57]. SigProfilerTopography allows performing analysis of strand-coordinated mutagenesis by identifying groups of consecutive mutated single base substitutions, attributed to the same mutational signatures, with no more than 10kb distance between any two mutations. Mutations are oriented by the reference base of the Watson-Crick base-pair to ensure that they are occurring on the same strand, *e.g.*, consecutive C>A mutations attributed to a single mutational signature. Groups of varying lengths are pooled across all samples for each mutational signature. Same procedure is repeated for simulated mutations to assess the statistical significance of the observed number of strand-coordinated mutagenesis groups with expected list of number of strand-coordinated mutagenesis groups for each group length (**Fig. 5*A-C***).

**Figure 5.**
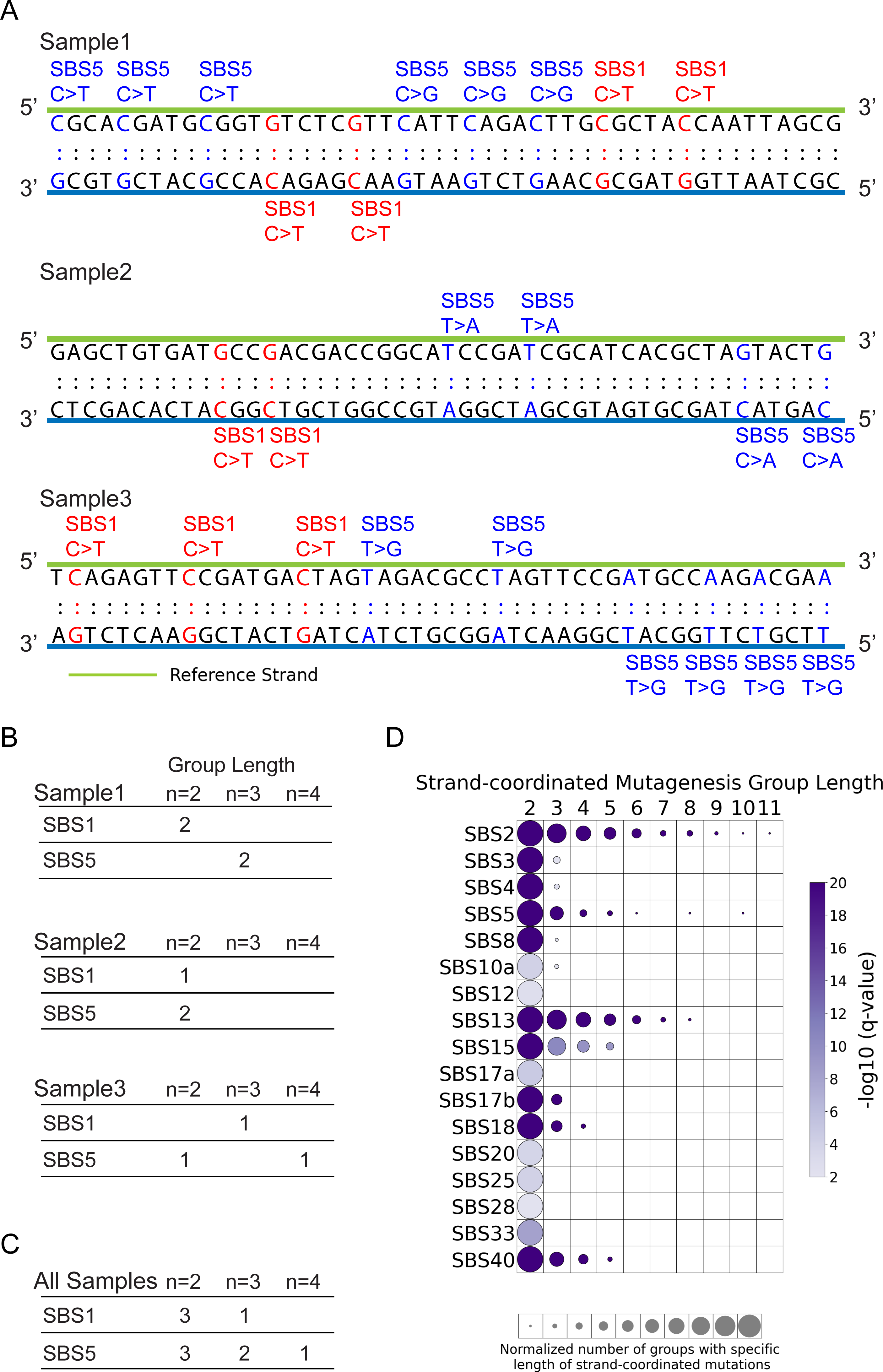
**Mapping strand-coordinated mutagenesis. *(A)*** Three simplified exemplar samples illustrating consecutive single base substitutions occurring on the same DNA strand due to specific mutational signatures. For example, consecutive three C>T mutations on the same strand generated by SBS5 within sample 1 result in one strand-coordinated mutagenesis group of length 3. ***(B)*** Summary of strand-coordinated mutagenesis groups of varying lengths for each mutational signature within each of the three examined samples from panel *(A)*. ***(C)*** Accumulation of strand-coordinated mutagenesis groups across all three examined exemplar samples from panel *(A)*. ***(D)*** Strand-coordinated mutagenesis for COSMIC substitution signatures operative in 552 ESCCs. Circle plot displays the group lengths from 2 to 11 mutations on the x-axis and the SBS mutational signatures on the y-axis. Circle size represents the number of strand-coordinated mutagenesis groups for the corresponding group length, which is normalized for each mutational signature. Circle color indicates the statistical significance of the finding with -log10 (q-value), with darker color corresponding to lower q-value.

Applying SigProfilerTopography to all mutational signatures operative in the 552 whole-genome sequenced samples revealed statistically significant strand-coordinated mutagenesis for multiple signatures. The APOBEC-attributed signatures SBS2 and SBS13 exhibited groups of up to 11 consecutive mutations likely due to APOBEC-induced kataegis [58, 59]. Interestingly, the flat signatures SBS5 and SBS40 also manifested strand-coordinated mutagenesis of varying group length. Lastly, the mismatch repair deficiency signature SBS15 also exhibited strand-coordinated mutagenesis for as many as 5 consecutively mutated bases (**Fig. 5*D***).

## DISCUSSION

SigProfilerTopography is an open-source Python package that allows understanding the interplay between somatic mutagenesis and the structural and topographical features of a genome. The tool can reveal mutational signature-specific tendencies associated with chromatin organization, histone modifications, and transcription factor binding as well as ones affected by cellular processes such as DNA replication and transcription. As we illustrated by applying the tool to 552 whole-genome sequenced ESCCs, SigProfilerTopography simultaneously examines real somatic mutations and simulated mutations, compares their tendencies, and then elucidates the statistically significant differences for each structural and topographical feature of interest. The tool also seamlessly integrates with other SigProfiler tools and leverages them for parts of its computational workflow, including classification of somatic mutations using SigProfilerMatrixGenerator [60], simulating realistic background mutations with SigProfilerSimulator [38], and assigning mutational signatures to each somatic mutation using SigProfilerAssignment [36].

SigProfilerTopography has at least three known limitations. First, the tool can only be used to explore small mutational events including single base substitutions, doublet substitutions, and small insertions and deletions. Currently, the tool does not allow exploring large mutational events [61] such as copy-number changes and structural rearrangements. Second, SigProfilerTopography can be applied only to whole-genome sequenced cancers, and it will not work on whole-exome or targeted cancer gene panel sequencing data as the algorithm requires profiling the non-coding regions of the genome. Lastly, the tool necessitates sufficient numbers of somatic mutations for the statistical analyses to be meaningful and statistically significant. We have previously shown that topographical analyses will work and can yield biologically exciting results when examining adult cancers [6], however, it is currently unclear whether some pediatric cancer genomes will have sufficient numbers of somatic mutations for examining the topography of their mutational signatures.

## CONCLUSIONS

SigProfilerTopography enables a thorough examination of how genome topography and genome architecture impact the accrual of somatic mutations. The tool offers a robust approach for evaluation of localized somatic mutation rates across various genomic features within a single comprehensive platform, offering a scalable solution for analyzing large datasets encompassing many thousands of cancer genomes and all types of small mutational event. Overall, SigProfilerTopography is a computational tool that provides an unprecedented opportunity for understanding the biological mechanisms and molecular processes influencing somatic mutational processes that have operated in a cancer genome.

## METHODS

### Tool implementation

SigProfilerTopography is developed as a computationally efficient Python package, and it is available for installation through PyPI. The tool leverages SigProfilerAssignment for attributing mutational signatures to individual somatic mutations [36], SigProfilerSimulator for generating all simulated datasets [38], and SigProfilerMatrixGenerator for processing input data for somatic mutations [60]. SigProfilerTopography allows processing all types of small mutational events, including: *(i)* single base substitutions, *(ii)* doublet base substitutions, and *(iii)* small insertions and deletions. The tool supports most commonly used data formats for somatic mutations: Variant Calling Format (VCF), Mutation Annotation Format (MAF), International Cancer Genome Consortium (ICGC) data format, and simple text file. SigProfilerTopography allows examining topography features in wiggle (wig), browser extensible data (bed), bigWig, and bigBed formats. The tool has been extensively tested on data from transposase-accessible chromatin with sequencing (ATAC-Seq), replication sequencing (Repli-Seq), micrococcal nuclease sequencing (MNase-Seq), and immunoprecipitation sequencing (ChIP-Seq). By default, the tool performs statistical comparisons and Benjamini-Hochberg corrections for multiple hypothesis testing using the statsmodels Python package. SigProfilerTopography is freely available, distributed under the BSD-2-Clause license, and has been extensively documented.

*Python code:* https://github.com/AlexandrovLab/SigProfilerTopography

*Documentation:* https://osf.io/5unby/wiki/home/

### Esophageal cancer dataset

A previous study [34] collected 552 esophageal squamous cell carcinomas (ESCC) including tumor and germline DNA, which were subjected to whole-genome sequencing with mean sequencing coverage of 49-fold and 26-fold, respectively. *De novo* mutational signatures were extracted and decomposed into COSMIC reference signatures using SigProfilerExtractor [33]. All somatic mutations within the ESCC dataset were considered with each mutation probabilistically assigned to each of the operative mutational signatures.

## ABBREVIATIONS

ATAC-Seq: assay for transposase-accessible chromatin with sequencing
Bed: browser extensible data
ChIP-Seq: chromatin immunoprecipitation sequencing
COSMIC: Catalogue Of Somatic Mutations In Cancer
DBS: doublet base substitutions
ENCODE: Encyclopedia of DNA Elements
ESCC: esophageal squamous cell carcinoma
ICGC: International Cancer Genome Consortium
ID: small insertions and deletions
Kb: kilobase
MAF: Mutation Annotation Format
MNase-Seq: micrococcal nuclease sequencing
Repli-Seq: replication sequencing
SBS: single base substitutions
VCF: Variant Call Format
Wig: wiggle

## DECLARATIONS

### Ethics approval and consent to participate

Not applicable.

### Consent for publication

Not applicable.

### Availability of data and materials

Data sharing is not applicable to this article as no datasets were generated during the current study. The somatic mutations for the 552 previously generated esophageal squamous cell carcinoma were retrieved from: https://doi.org/10.6084/m9.figshare.22744733.

### Authors’ contributions

BO developed the Python code and wrote the draft of the manuscript. LBA supervised the overall development of the code and writing of the manuscript. All authors read and approved the final manuscript.

## Acknowledgements

The computational development reported in this manuscript have utilized the Triton Shared Computing Cluster at the San Diego Supercomputer Center of UC San Diego. We thank Marcos Díaz-Gay for reading the draft of the manuscript and for providing feedback. We also thank Ting Yang, Mousumy Kundu, and Mark Barnes for testing the tool and for the helpful discussions. The research in this study was also supported by UC San Diego Sanford Stem Cell Institute.

## REFERENCES

1. Martincorena I, Campbell PJ: **Somatic mutation in cancer and normal cells**. Science 2015, 349:1483–1489.

2. Consortium ITP-CAoWG: **Pan-cancer analysis of whole genomes**. Nature 2020, 578:82–93.

3. Alexandrov LB, Nik-Zainal S, Wedge DC, Campbell PJ, Stratton MR: **Deciphering signatures of mutational processes operative in human cancer**. Cell Rep 2013, 3:246–259.

4. Alexandrov LB, Stratton MR: Mutational signatures: the patterns of somatic mutations hidden in cancer genomes. Curr Opin Genet Dev 2014, 24:52–60.

5. Alexandrov LB, Kim J, Haradhvala NJ, Huang MN, Tian Ng AW, Wu Y, Boot A, Covington KR, Gordenin DA, Bergstrom EN, et al: The repertoire of mutational signatures in human cancer. Nature 2020, 578:94–101.

6. Otlu B, Diaz-Gay M, Vermes I, Bergstrom EN, Zhivagui M, Barnes M, Alexandrov LB: **Topography of mutational signatures in human cancer**. Cell Rep 2023, 42:112930.

7. Schuster-Bockler B, Lehner B: Chromatin organization is a major influence on regional mutation rates in human cancer cells. Nature 2012, 488:504–507.

8. Tomkova M, Tomek J, Kriaucionis S, Schuster-Bockler B: Mutational signature distribution varies with DNA replication timing and strand asymmetry. Genome Biol 2018, 19:129.

9. Stamatoyannopoulos JA, Adzhubei I, Thurman RE, Kryukov GV, Mirkin SM, Sunyaev SR: **Human mutation rate associated with DNA replication timing**. Nat Genet 2009, 41:393–395.

10. Frigola J, Sabarinathan R, Mularoni L, Muinos F, Gonzalez-Perez A, Lopez-Bigas N: **Reduced mutation rate in exons due to differential mismatch repair**. Nat Genet 2017, 49:1684–1692.

11. Imielinski M, Guo G, Meyerson M: Insertions and Deletions Target Lineage-Defining Genes in Human Cancers. Cell 2017, 168:460–472 e414.

12. Brown AJ, Mao P, Smerdon MJ, Wyrick JJ, Roberts SA: **Nucleosome positions establish an extended mutation signature in melanoma**. PLoS Genet 2018, 14:e1007823.

13. Pich O, Muinos F, Sabarinathan R, Reyes-Salazar I, Gonzalez-Perez A, Lopez-Bigas N: **Somatic and Germline Mutation Periodicity Follow the Orientation of the DNA Minor Groove around Nucleosomes**. Cell 2018, 175:1074–1087 e1018.

14. Gonzalez-Perez A, Sabarinathan R, Lopez-Bigas N: **Local Determinants of the Mutational Landscape of the Human Genome**. Cell 2019, 177:101–114.

15. Li F, Mao G, Tong D, Huang J, Gu L, Yang W, Li GM: The histone mark H3K36me3 regulates human DNA mismatch repair through its interaction with MutSalpha. Cell 2013, 153:590–600.

16. Sabarinathan R, Mularoni L, Deu-Pons J, Gonzalez-Perez A, Lopez-Bigas N: **Nucleotide excision repair is impaired by binding of transcription factors to DNA**. Nature 2016, 532:264–267.

17. Katainen R, Dave K, Pitkanen E, Palin K, Kivioja T, Valimaki N, Gylfe AE, Ristolainen H, Hanninen UA, Cajuso T, et al: CTCF/cohesin-binding sites are frequently mutated in cancer. Nat Genet 2015, 47:818–821.

18. Morganella S, Alexandrov LB, Glodzik D, Zou X, Davies H, Staaf J, Sieuwerts AM, Brinkman AB, Martin S, Ramakrishna M, et al: The topography of mutational processes in breast cancer genomes. Nat Commun 2016, 7:11383.

19. Haradhvala NJ, Polak P, Stojanov P, Covington KR, Shinbrot E, Hess JM, Rheinbay E, Kim J, Maruvka YE, Braunstein LZ, et al: Mutational Strand Asymmetries in Cancer Genomes Reveal Mechanisms of DNA Damage and Repair. Cell 2016, 164:538–549.

20. Fischer A, Illingworth CJ, Campbell PJ, Mustonen V: EMu: probabilistic inference of mutational processes and their localization in the cancer genome. Genome Biol 2013, 14:R39.

21. Mayakonda A, Lin DC, Assenov Y, Plass C, Koeffler HP: **Maftools: efficient and comprehensive analysis of somatic variants in cancer**. Genome Research 2018, 28:1747–1756.

22. Blokzijl F, Janssen R, van Boxtel R, Cuppen E: MutationalPatterns: comprehensive genome-wide analysis of mutational processes. Genome Med 2018, 10:33.

23. Fantini D, Vidimar V, Yu YN, Condello S, Meeks JJ: MutSignatures: an R package for extraction and analysis of cancer mutational signatures. Scientific Reports 2020, 10.

24. Ardin M, Cahais V, Castells X, Bouaoun L, Byrnes G, Herceg Z, Zavadil J, Olivier M: **MutSpec: a Galaxy toolbox for streamlined analyses of somatic mutation spectra in human and mouse cancer genomes**. BMC Bioinformatics 2016, 17:170.

25. Gori K, Baez-Ortega A: sigfit: flexible Bayesian inference of mutational signatures. *bioRxiv* 2020:372896.

26. Wang S, Li H, Song M, Tao Z, Wu T, He Z, Zhao X, Wu K, Liu XS: Copy number signature analysis tool and its application in prostate cancer reveals distinct mutational processes and clinical outcomes. PLoS Genet 2021, 17:e1009557.

27. Kasar S, Kim J, Improgo R, Tiao G, Polak P, Haradhvala N, Lawrence MS, Kiezun A, Fernandes SM, Bahl S, et al: Whole-genome sequencing reveals activation-induced cytidine deaminase signatures during indolent chronic lymphocytic leukaemia evolution. Nat Commun 2015, 6:8866.

28. Degasperi A, Amarante TD, Czarnecki J, Shooter S, Zou XQ, Glodzik D, Morganella S, Nanda AS, Badja C, Koh G, et al: A practical framework and online tool for mutational signature analyses show intertissue variation and driver dependencies (vol 1, pg 249, 2020). Nature Cancer 2020, 1:748-748.

29. Rosales RA, Drummond RD, Valieris R, Dias-Neto E, da Silva IT: **signeR: an empirical Bayesian approach to mutational signature discovery**. Bioinformatics 2017, 33:8–16.

30. Gehring JS, Fischer B, Lawrence M, Huber W: SomaticSignatures: inferring mutational signatures from single-nucleotide variants. Bioinformatics 2015, 31:3673–3675.

31. Vohringer H, Hoeck AV, Cuppen E, Gerstung M: Learning mutational signatures and their multidimensional genomic properties with TensorSignatures. Nat Commun 2021, 12:3628.

32. Lee J, Lee AJ, Lee JK, Park J, Kwon Y, Park S, Chun H, Ju YS, Hong D: Mutalisk: a web-based somatic MUTation AnaLyIS toolKit for genomic, transcriptional and epigenomic signatures. Nucleic Acids Res 2018, 46:W102–W108.

33. Islam SMA, Diaz-Gay M, Wu Y, Barnes M, Vangara R, Bergstrom EN, He Y, Vella M, Wang J, Teague JW, et al: Uncovering novel mutational signatures by de novo extraction with SigProfilerExtractor. Cell Genom 2022, 2.

34. Moody S, Senkin S, Islam SMA, Wang J, Nasrollahzadeh D, Cortez Cardoso Penha R, Fitzgerald S, Bergstrom EN, Atkins J, He Y, et al: **Mutational signatures in esophageal squamous cell carcinoma from eight countries with varying incidence**. Nat Genet 2021, 53:1553–1563.

35. Consortium EP: An integrated encyclopedia of DNA elements in the human genome. Nature 2012, 489:57-74.

36. Diaz-Gay M, Vangara R, Barnes M, Wang X, Islam SMA, Vermes I, Duke S, Narasimman NB, Yang T, Jiang Z, et al: Assigning mutational signatures to individual samples and individual somatic mutations with SigProfilerAssignment. Bioinformatics 2023, 39.

37. Tate JG, Bamford S, Jubb HC, Sondka Z, Beare DM, Bindal N, Boutselakis H, Cole CG, Creatore C, Dawson E, et al: COSMIC: the Catalogue Of Somatic Mutations In Cancer. Nucleic Acids Res 2019, 47:D941–D947.

38. Bergstrom EN, Barnes M, Martincorena I, Alexandrov LB: Generating realistic null hypothesis of cancer mutational landscapes using SigProfilerSimulator. BMC Bioinformatics 2020, 21:438.

39. Koh G, Degasperi A, Zou X, Momen S, Nik-Zainal S: **Mutational signatures: emerging concepts, caveats and clinical applications**. Nat Rev Cancer 2021, 21:619–637.

40. Christensen S, Van der Roest B, Besselink N, Janssen R, Boymans S, Martens JWM, Yaspo ML, Priestley P, Kuijk E, Cuppen E, Van Hoeck A: **5-Fluorouracil treatment induces characteristic T>G mutations in human cancer**. Nat Commun 2019, 10:4571.

41. Zhou C, Greenberg MM: DNA damage by histone radicals in nucleosome core particles. J Am Chem Soc 2014, 136:6562–6565.

42. Hara R, Mo JY, Sancar A: DNA damage in the nucleosome core is refractory to repair by human excision nuclease. Molecular and Cellular Biology 2000, 20:9173–9181.

43. Calo E, Wysocka J: Modification of Enhancer Chromatin: What, How, and Why? Molecular Cell 2013, 49:825–837.

44. Kang Y, Kim YW, Kang J, Kim A: Histone H3K4me1 and H3K27ac play roles in nucleosome eviction and eRNA transcription, respectively, at enhancers. FASEB J 2021, 35:e21781.

45. Marchal C, Sasaki T, Vera D, Wilson K, Sima J, Rivera-Mulia JC, Trevilla-Garcia C, Nogues C, Nafie E, Gilbert DM: **Genome-wide analysis of replication timing by next-generation sequencing with E/L Repli-seq**. Nat Protoc 2018, 13:819–839.

46. Gilbert DM: Making sense of eukaryotic DNA replication origins. Science 2001, 294:96-100.

47. Ryba T, Battaglia D, Pope BD, Hiratani I, Gilbert DM: **Genome-scale analysis of replication timing: from bench to bioinformatics**. Nat Protoc 2011, 6:870–895.

48. Hansen RS, Thomas S, Sandstrom R, Canfield TK, Thurman RE, Weaver M, Dorschner MO, Gartler SM, Stamatoyannopoulos JA: **Sequencing newly replicated DNA reveals widespread plasticity in human replication timing**. Proc Natl Acad Sci U S A 2010, 107:139–144.

49. Thurman RE, Day N, Noble WS, Stamatoyannopoulos JA: Identification of higher-order functional domains in the human ENCODE regions. Genome Res 2007, 17:917–927.

50. Bell SP, Dutta A: **DNA replication in eukaryotic cells**. Annu Rev Biochem 2002, 71:333–374.

51. Hoopes JI, Cortez LM, Mertz TM, Malc EP, Mieczkowski PA, Roberts SA: APOBEC3A and APOBEC3B Preferentially Deaminate the Lagging Strand Template during DNA Replication. Cell Rep 2016, 14:1273–1282.

52. Sancar A: Mechanisms of DNA Repair by Photolyase and Excision Nuclease (Nobel Lecture). Angew Chem Int Ed Engl 2016, 55:8502-8527.

53. Hanawalt PC, Spivak G: Transcription-coupled DNA repair: two decades of progress and surprises. Nat Rev Mol Cell Biol 2008, 9:958–970.

54. Fujimoto A, Furuta M, Totoki Y, Tsunoda T, Kato M, Shiraishi Y, Tanaka H, Taniguchi H, Kawakami Y, Ueno M, et al: Whole-genome mutational landscape and characterization of noncoding and structural mutations in liver cancer. Nat Genet 2016, 48:500–509.

55. Letouze E, Shinde J, Renault V, Couchy G, Blanc JF, Tubacher E, Bayard Q, Bacq D, Meyer V, Semhoun J, et al: Mutational signatures reveal the dynamic interplay of risk factors and cellular processes during liver tumorigenesis. Nat Commun 2017, 8:1315.

56. Saini N, Gordenin DA: **Hypermutation in single-stranded DNA**. DNA Repair (Amst) 2020, 91-92:102868.

57. Roberts SA, Sterling J, Thompson C, Harris S, Mav D, Shah R, Klimczak LJ, Kryukov GV, Malc E, Mieczkowski PA, et al: Clustered mutations in yeast and in human cancers can arise from damaged long single-strand DNA regions. Mol Cell 2012, 46:424–435.

58. Nik-Zainal S, Alexandrov LB, Wedge DC, Van Loo P, Greenman CD, Raine K, Jones D, Hinton J, Marshall J, Stebbings LA, et al: Mutational processes molding the genomes of 21 breast cancers. Cell 2012, 149:979–993.

59. Roberts SA, Lawrence MS, Klimczak LJ, Grimm SA, Fargo D, Stojanov P, Kiezun A, Kryukov GV, Carter SL, Saksena G, et al: An APOBEC cytidine deaminase mutagenesis pattern is widespread in human cancers. Nature Genetics 2013, 45:970-+.

60. Bergstrom EN, Huang MN, Mahto U, Barnes M, Stratton MR, Rozen SG, Alexandrov LB: **SigProfilerMatrixGenerator: a tool for visualizing and exploring patterns of small mutational events**. BMC Genomics 2019, 20:685.

61. Khandekar A, Vangara R, Barnes M, Diaz-Gay M, Abbasi A, Bergstrom EN, Steele CD, Pillay N, Alexandrov LB: **Visualizing and exploring patterns of large mutational events with SigProfilerMatrixGenerator**. BMC Genomics 2023, 24:469.

